# Enhancing gRNA Transcript levels by Reducing the Scaffold Poly-T Tract for Optimal SpCas9- and SaCas9-mediated Gene Editing

**DOI:** 10.1101/2024.07.19.604224

**Authors:** Yu C. J. Chey, Luke Gierus, Caleb Lushington, Jayshen C. Arudkumar, Ashleigh Geiger, Lachlan G. Staker, Louise J. Robertson, Chandran Pfitzner, Jesse G. Kennedy, Ryan H. B. Lee, Gelshan I. Godahewa, Paul Q. Thomas, Fatwa Adikusuma

## Abstract

Ensuring sufficient gRNA transcript levels is critical for obtaining optimal CRISPR-Cas9 gene editing efficiency. The standard gRNA scaffold contains a sequence of four thymine nucleotides (4T), which is known to inhibit transcription from Pol III promoters such as the U6 promoter. Our study showed that using standard plasmid transfection protocols, the presence of these 4Ts did not significantly affect editing efficiency, as most of the gRNAs tested (55 gRNAs) achieved near-perfect editing outcomes. We observed that gRNAs with lower activity were T-rich and had reduced gRNA transcript levels. However, this issue can be effectively resolved by increasing transcript levels, which can be readily achieved by shortening the 4T sequences. In this study, we demonstrated this by modifying the sequences to 3TC. Although the 3TC scaffold modification did not improve editing efficiency for already efficient gRNAs when high vector quantities were available, it proved highly beneficial under conditions of limited vector availability, where the 3TC scaffold yielded higher editing efficiency. Additionally, we demonstrated that the 3TC scaffold is compatible with SpCas9 high-fidelity variants and ABEmax base editing, enhancing their editing efficiency. Another commonly used natural Cas9 variant, SaCas9, also benefited from the 3TC scaffold sequence modification, which increased gRNA transcription and subsequently improved editing activity. This modification was applied to the EDIT-101 therapeutic strategy, where it demonstrated marked improvements in performance. This study highlights the importance of shortening the 4T sequences in the gRNA scaffold to optimize gRNA transcript expression for enhanced CRISPR-Cas9 gene editing efficiency. This optimization is particularly important for therapeutic applications, where the quantity of vector is often limited, ensuring more effective and optimal outcomes.

## Introduction

Clustered Regularly Interspaced Short Palindromic Repeats/CRISPR-associated protein 9 (CRISPR-Cas9) gene-editing technology is a simple yet powerful tool for the targeted modification of DNA. This technology has been widely adopted for fundamental research applications and the development of precision therapeutics for genetic diseases. Originally derived from the anti-viral defence system of prokaryotes, CRISPR-Cas9 has been refined into a two-component ribonucleoprotein system, consisting of (i) a Cas9 endonuclease that binds to a protospacer adjacent motif (PAM) sequence and (ii) a programmable single-stranded guide RNA (gRNA) that confers target specificity (Barrangou and Doudna, 2016, Jinek et al., 2012, Doudna and Charpentier, 2014, Hsu et al., 2014). Within the cell nucleus, the Cas9-gRNA complex binds to PAM-containing DNA, creating a double-stranded break if the upstream sequence matches the gRNA target sequence. The DNA break prompts the endogenous DNA repair machinery to repair the free ends through template-independent non-homologous end joining (NHEJ) or template-dependent homology-directed repair pathways. In the case of NHEJ repair, the target DNA is prone to be repaired with errors resulting in sequence insertions or deletions (InDel) (Cong et al., 2013, Mali et al., 2013, Rouet et al., 1994).

The CRISPR genome editing toolkit includes a variety of natural and engineered Cas9 variants. The most widely used natural Cas9, *Streptococcus pyogenes* Cas9 (SpCas9), recognises and binds to the canonical NGG PAM (Cong et al., 2013, Jinek et al., 2012, Mali et al., 2013). Engineered variants of SpCas9, such as SpCas9-HF1 (Kleinstiver et al., 2016) and eSpCas9(1.1) (Slaymaker et al., 2016), have been developed to improve target specificity and mitigate off-target activity (Fu et al., 2013). The extended toolkit also includes Cas9- fusion proteins such as adenine and cytosine base editors as well as Prime editors (Anzalone et al., 2019, Gaudelli et al., 2017, Komor et al., 2016). For example, adenine base editors (ABEs) utilise the Cas9D10A nickase fused to a deoxyadenosine deaminase to enable DNA nucleotide conversions of A•T to G•C (Gaudelli et al., 2017), and engineered ABEs such as the ABEmax have been optimised for reduced RNA off-targeting (Rees and Liu, 2018, Li et al., 2021). *Staphylococcus aureus* Cas9 (SaCas9), another natural Cas9 ortholog, recognises the NNGRRT PAM and has practical applications for in-vivo genome editing due to its relatively small size, allowing for complete packaging in a single adenoviral-associated virus (AAV) vector (Ran et al., 2015).

Achieving high editing efficiency is pivotal for the diverse applications of CRISPR gene-editing technology. Enhanced efficacies of CRISPR-based gene editing can lead to improved therapeutic outcomes, increased efficiency in generating genetic models, and the development of more powerful tools such as for gene drive applications. Ensuring the sufficient availability of gRNA transcripts is a key factor in attaining optimal editing efficiency. In this study, we demonstrated that despite the presence of a 4T sequence known to hinder Pol III transcription in the SpCas9 gRNA scaffold (Arimbasseri and Maraia, 2015, Gao et al., 2018), we achieved near-perfect editing efficiency for most of the gRNAs tested using standard transfection protocols. Furthermore, we found that shortening the 4T sequences significantly increased gRNA expression, effectively overcoming suboptimal editing efficiency caused by lower gRNA transcript levels. This optimized gRNA expression proved to be beneficial for various gene editing applications, including those utilizing SpCas9 high-fidelity variants, base editing, SaCas9, and scenarios where vector availability is limited.

## Results

### SpCas9 with original gRNA scaffold can perform genome editing with near-perfect editing efficiency

Using the all-in-one CRISPR plasmid PX459.v2 (CBh-driven SpCas9-T2A-Puro and U6- driven single gRNA) with the original gRNA scaffold containing a sequence of 4 thymine nucleotides (4T) (Ran et al., 2013), we tested 55 different SpCas9 gRNAs across several target genes. Experiments were conducted in three different cell lines by plasmid transfection with subsequent puromycin selection. These target genes were: *EMX1, VEGFA, RUNX1* and *DMD* in human HEK293T cells; *Dmd* in mouse C2C12 cells; and *PolQ1, 53bp1, Mixl1* and *Prl* in mouse embryonic stem (mES) cells. Despite the inhibitory effect of the 4T sequence in the gRNA scaffold on Pol III promoter transcription (Gao et al., 2018, Arimbasseri and Maraia, 2015), most of the gRNAs tested displayed near-perfect editing, with ten gRNAs tested in HEK293T generating over 95% modified reads and only one gRNA (hDMD-B) with slightly lower editing at ∼92.0% (Figure 1A). In C2C12 cells, 12 out of 16 gRNAs generated over 90% modified alleles, while the remaining four gRNAs (mDmd-E, -H, -I and -K) had moderate editing efficiency of around 83-89% modified alleles (Figure 1B). In mES cells, all 27 gRNAs had around 99% editing efficiency, except for the mPrl 1.4 gRNA, which had a slightly lower efficiency of 93.6% (Figure 1C).

**Figure 1:**
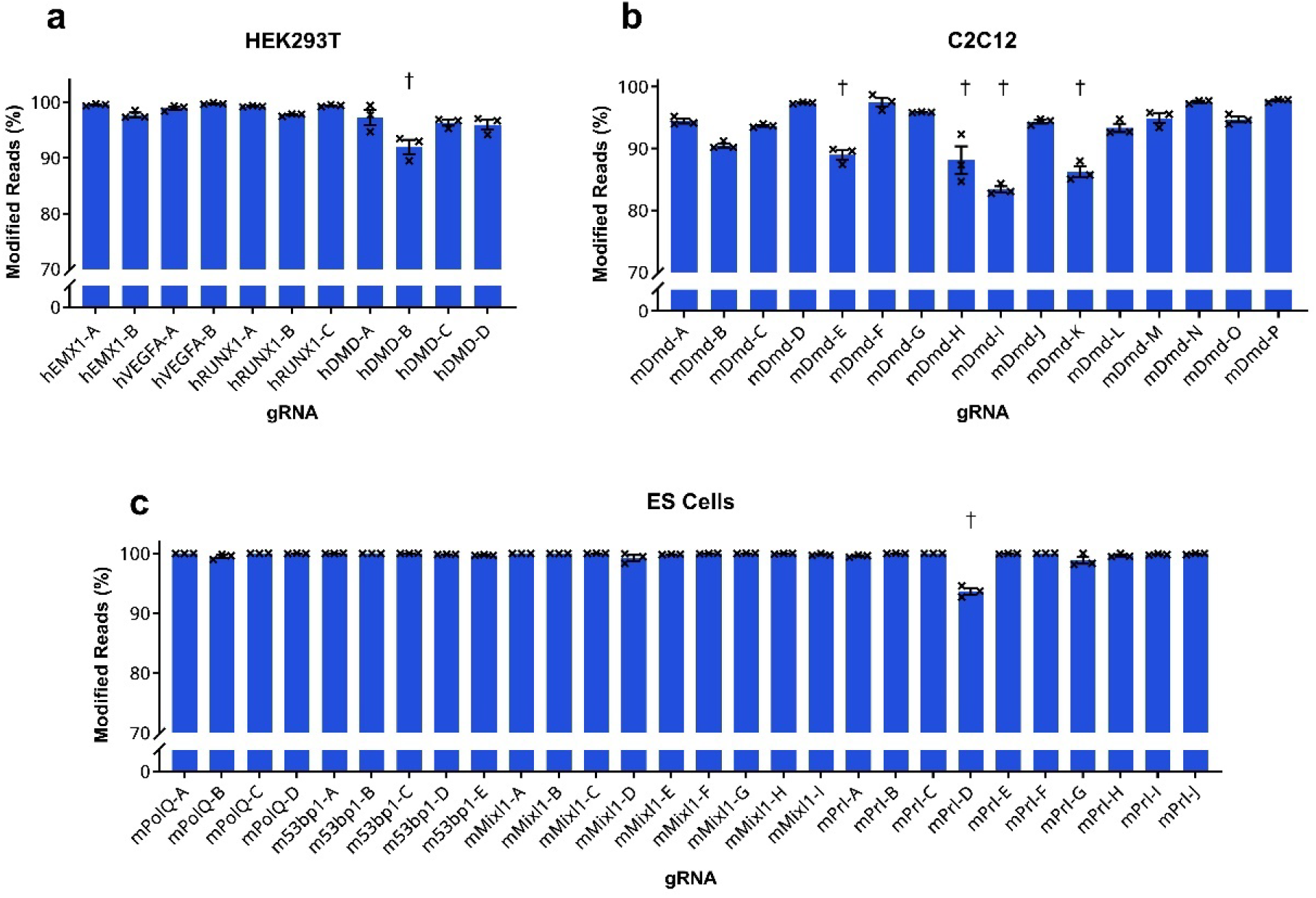
Assessment of editing efficiencies of original (4T) scaffold Sp gRNAs by deep amplicon sequencing in (A) HEK293T, (B) C2C12 and (C) mES cells. PX459.V2 plasmids were delivered via nucleofection with subsequent application of puromycin selection. Mean ± SEM, n=3. Obelisks (†) indicate gRNAs with lower editing efficiencies compared to the performance of other gRNAs tested in the same cell line.

### Addressing lower editing efficiency through increased gRNA expression

We hypothesised that the comparatively lower editing efficiencies exhibited by mDmd-E, -H, -I, -K and mPrl-D could be attributed to lower gRNA transcript levels produced from the vectors. These gRNAs are generally T-rich in sequence, and the presence of 4Ts in the gRNA scaffold likely exacerbates this issue by negatively impacting RNA pol III transcription (Graf et al., 2019, DeWeirdt et al., 2022). To investigate this, we measured the gRNA transcript expression levels of three less-efficient gRNAs (mDmd-E, -H and -I) by qPCR and found that their gRNA transcript levels were lower than a control gRNA, mDmd-M (Figure 2B), consistent with our hypothesis.

**Figure 2:**
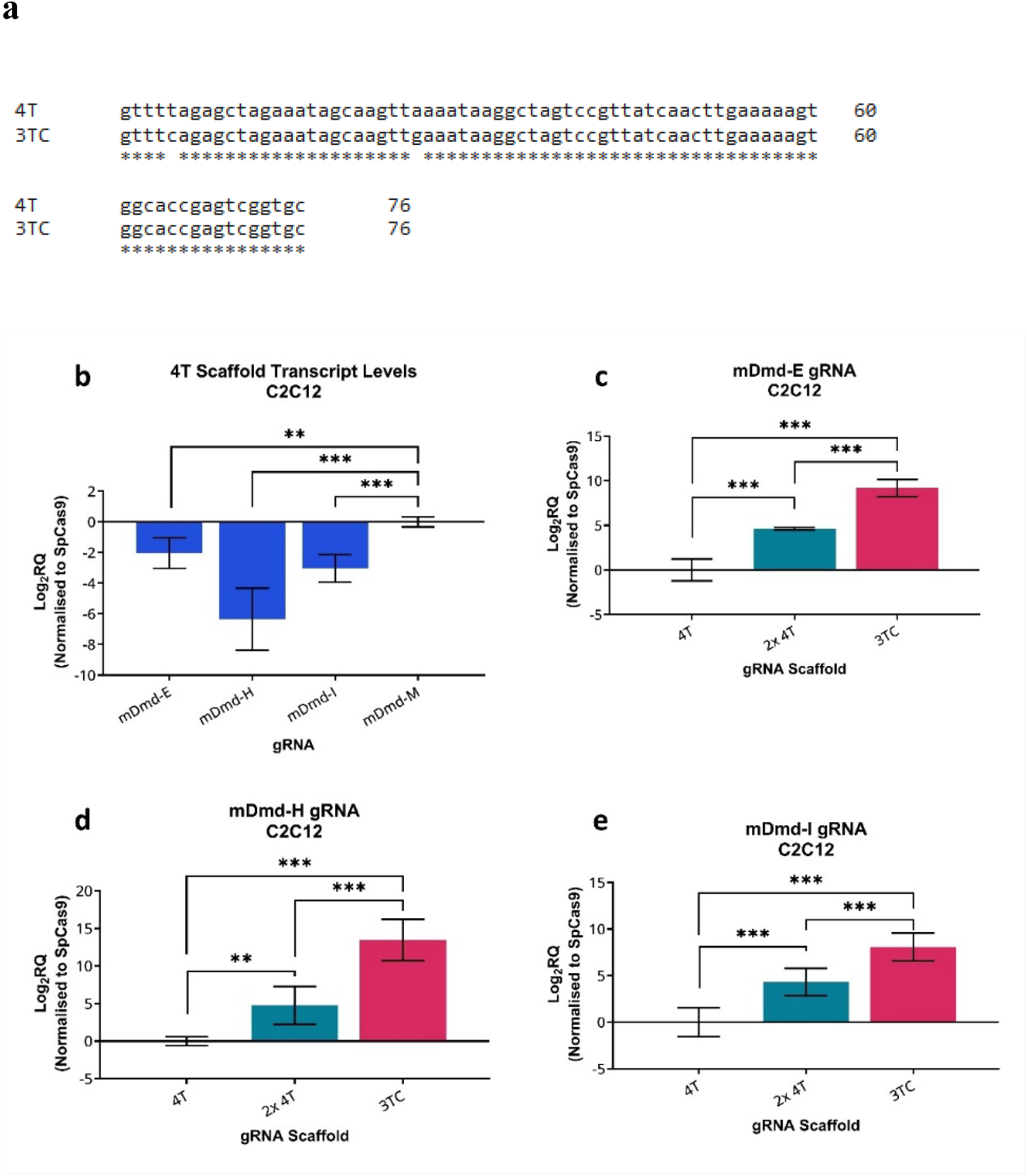
The modified 3TC scaffold boosts SpCas9 gRNA expression levels, compared to the original 4T scaffold. (A) DNA sequence of the 4T and modified 3TC scaffolds. (B) Relative quantification (RQ) of mDmd gRNA delivered by nucleofection of PX459.V2 (4T) to C2C12 cells, by qRT-PCR. Mean Log_2_RQ ± 95% CI; n=3. One-way ANOVA with Dunnett’s multiple comparisons test performed on ΔΔCT values, **p ≤ 0.01, ***p ≤ 0.001. (C, D, E) Comparison of the relative quantities of mDmd Sp gRNA, delivered by PX459.V2, pdg459.V2 (2x4T) and PX459.V3 (3TC), measured by qRT-PCR. Mean Log_2_RQ ± 95% CI; n=3. One-way ANOVA with Tukey’s multiple comparisons test performed on ΔΔCT values, **p ≤ 0.01, ***p ≤ 0.001.

We also hypothesised that increasing the gRNA transcript levels could improve the editing efficiency of these moderately-performing gRNAs. We tested two approaches to increase gRNA transcript levels. In the first, we doubled the U6 expression cassettes expressing a given gRNA using pDG459, our previously published dual-gRNA plasmid that was modified from PX459.V2 (Adikusuma et al., 2017). In the second approach, we performed a simple modification of the gRNA scaffold in the PX459.V2 plasmid to shorten the 4T-string by replacing the fourth T nucleotide in the tetraloop with a C nucleotide (3TC scaffold) and replacing its corresponding complementary A nucleotide with a G nucleotide (Figure 2A) (Dang et al., 2015, Gao et al., 2019).

Both methods significantly increased transcript expression of gRNA mDmd-E, -H and -I. While double-dosing with pDG459 increased gRNA levels, the single U6-driven PX459.V2 with 3TC scaffold appeared to be more effective and improved gRNA levels by a remarkable 8.1-13.5 doublings (271-11349 in fold changes) (Figure 2C-E). Following this, we tested the editing efficiencies of all lower efficient gRNAs with the dual-dosing and the 3TC scaffold approaches and found that their editing was enhanced to over 90% and over 95%, respectively (Figure 3A, B). The hDMD-B gRNA which had moderately high editing in HEK293Ts (∼92.0%) also had its editing boosted to ∼97.4% with the 3TC scaffold (Figure 3C). These findings indicate that the transcription levels of gRNA might be the critical limiting factor for the previously observed suboptimal editing efficiencies, which can be addressed by enhancing gRNA transcript levels. Furthermore, our data demonstrate that reducing the T-string in the gRNA scaffold, such as by employing the 3TC scaffold modification, is a straightforward and potent strategy to boost gRNA expression. This optimization is particularly beneficial for gRNAs with inherently lower efficiency due to reduced gRNA transcript levels, thereby achieving optimal editing performance.

**Figure 3:**
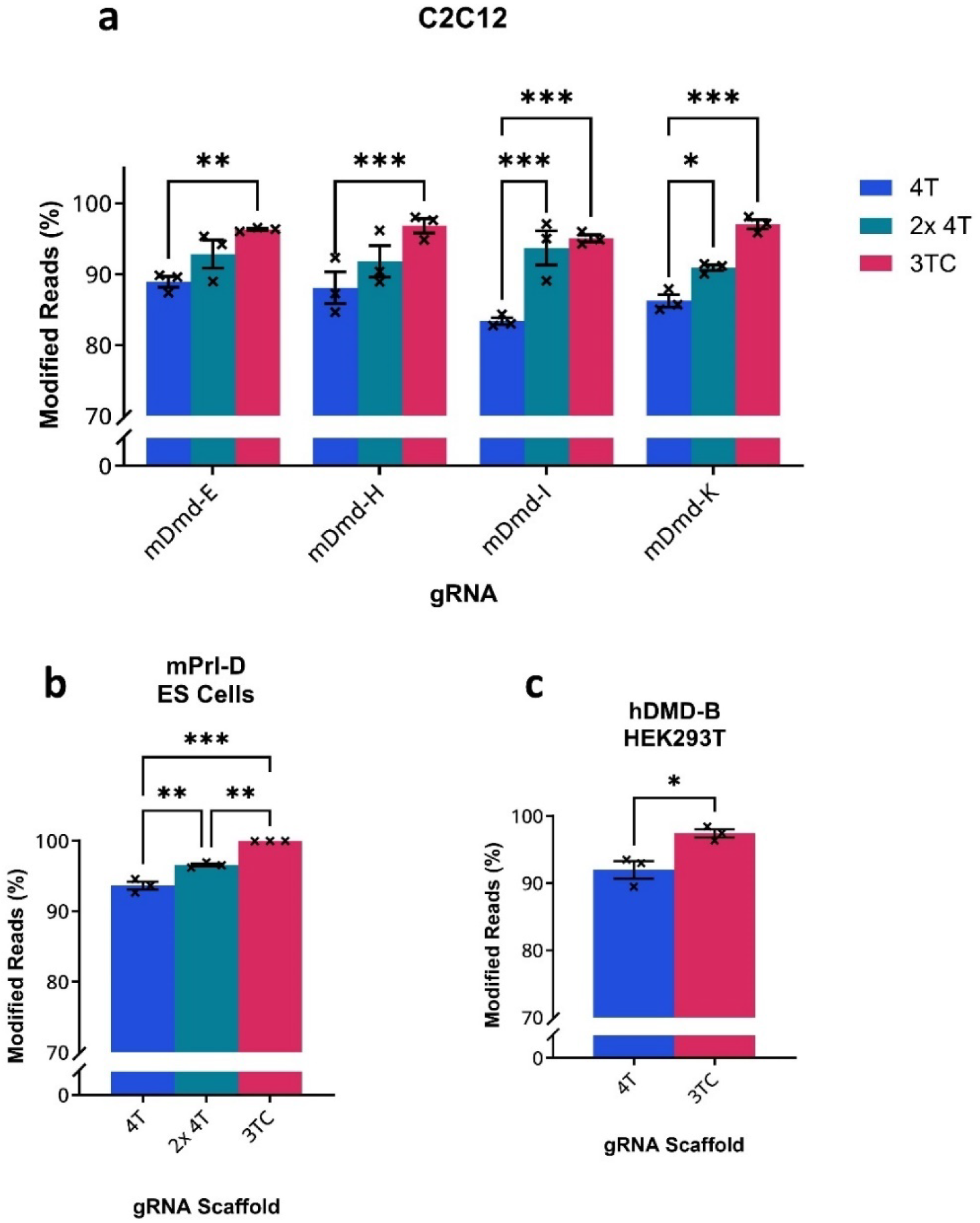
Editing efficiencies of 4T and 3TC modified scaffold gRNAs, assessed by deep amplicon sequencing. Comparison of nucleofected PX459.V2 (4T), pdg459.V2 (2x 4T) and PX459.V3 (3TC) plasmids on (A) *mDmd* gene editing in C2C12 cells and (B) mPrl-D editing in mES cells. Mean ± SEM; n=3. Statistical analysis performed by two-way ANOVA with Dunnett’s multiple comparisons test and one-way ANOVA with Tukey’s multiple comparisons, respectively. *p ≤ 0.05, **p ≤ 0.01, ***p ≤ 0.001. (C) Comparison of nucleofected PX459.V2 (4T) and PX459.V3 (3TC) plasmids on hDMD-B editing in HEK293T cells. Unpaired t-test, *p ≤ 0.05.

### Editing with low vector availability can benefit from enhanced transcript expression

In our previous experiments using the 4T scaffold, most gRNAs we tested demonstrated near-perfect editing efficiencies, suggesting that under standard transfection protocols, the conventional 4T scaffold is generally sufficient for achieving high editing efficiency. We hypothesised that the high efficiencies observed with the 4T scaffold may be attributed to the abundant vector quantity resulting from the transfection of high plasmid doses and subsequent transfectant selection. To explore this further, we conducted a comparison under conditions where the vector quantity was limited, by omitting the selection step. We observed that high levels of plasmid transfected without selection resulted in similar editing rates for the 3TC and 4T scaffolds. However, when vector quantity was restricted to a very low amount during transfection, we observed significantly higher editing efficiencies with the 3TC scaffold compared to the 4T scaffold (Figure 4A, B). This highlights the importance of enhanced transcript expression facilitated by the 3TC scaffold under conditions of limited vector availability.

**Figure 4:**
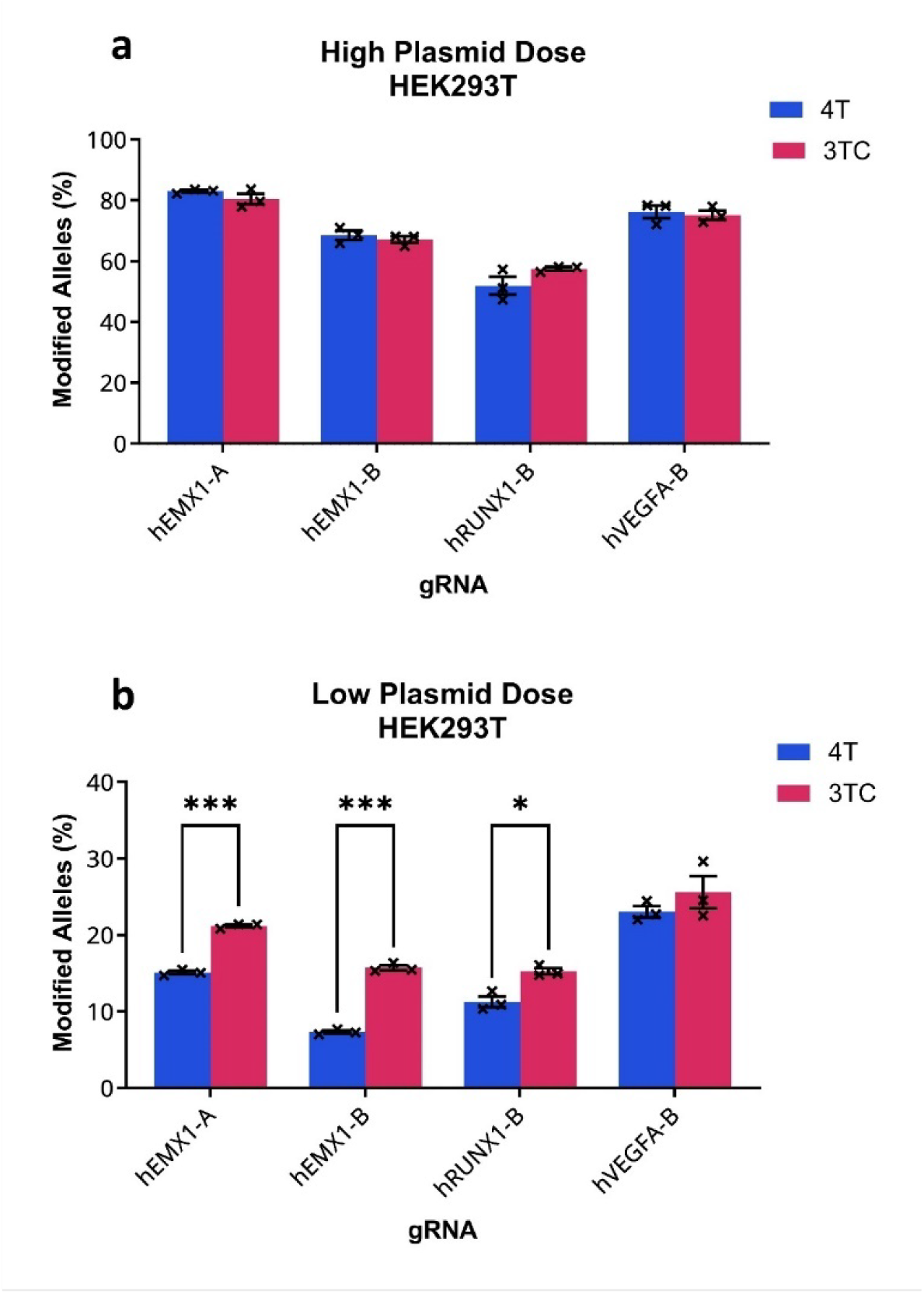
Editing efficiencies of PX459.V2 (4T) and PX459.V3 (3TC) in (A) high and (B) low plasmid doses delivered by lipofection without puromycin selection in HEK239T cells, assessed by deep amplicon sequencing. Mean ± SEM; n=3. Two-way ANOVA with Šídák’s multiple comparisons test; *p ≤ 0.05, ***p ≤ 0.001.

### The 3TC gRNA scaffold improves high-fidelity SpCas9s editing

Engineered high-fidelity SpCas9 variants, such as SpCas9-HF1 and eSpCas9(1.1), have been developed to enhance specificity and to reduce off-target activity. We aimed to investigate whether the 3TC scaffold modification is suitable and advantageous for these high-fidelity SpCas9 variants. To compare the performance of the 4T and 3TC scaffolds with these high-fidelity SpCas9 variants, we utilised all-in-one plasmids PX459.V2 SpCas9-HF1 and PX459.V2 eSpCas9(1.1) with the original 4T scaffold (Kato-Inui et al., 2018) to generate cognate 3TC versions which we denote as PX459.V3 SpCas9-HF1 and PX459.V3 eSpCas9(1.1). Four different gRNA spacers were assessed using high and low doses in HEK293T cells. Remarkably, we observed that the 3TC scaffold was not only compatible with SpCas9-HF1 and eSpCas9(1.1) but also enhanced their editing efficiency across various gRNAs, regardless of vector availability (Figure 5A, B).

**Figure 5:**
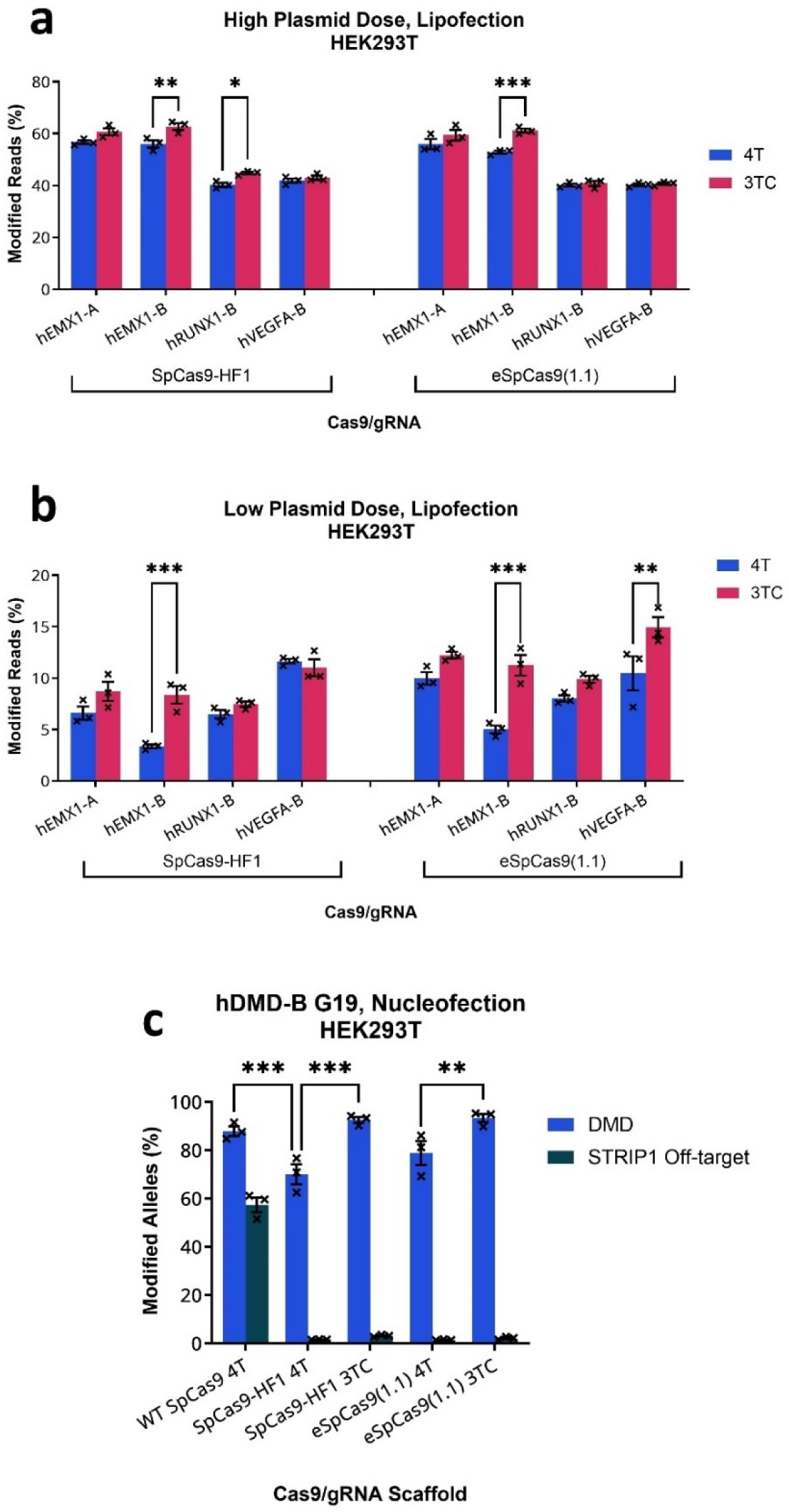
Editing efficiencies of high-fidelity SpCas9s with the 3TC scaffold. Comparison of PX459.V2 SpCas9-HF1 (4T), PX459.V3 SpCas9-HF1 (3TC), PX459.V2 eSpCas9(1.1) (4T) and PX459.V3 eSpCas9(1.1) (3TC) plasmids delivered by lipofection at a (A) high and (B) low plasmid dose without puromycin selection in HEK239T cells, assessed by deep amplicon sequencing. Mean ± SEM; n=3. Two-way ANOVA with Šídák’s multiple comparisons test; *p ≤ 0.05, **p ≤ 0.01, ***p ≤ 0.001. (C) Editing efficiencies of hDMD-B in the G19 gRNA configuration with WT and high-fidelity Sp-Cas9 plasmids delivered by nucleofection with puromycin selection in HEK293Ts. Two-way ANOVA with Šídák’s multiple comparisons test; *p ≤ 0.05, ***p ≤ 0.001.

We utilised this improvement for the gRNA, h51B, targeting exon-51 of the *DMD* gene, for single-cut editing to potentially reframe *DMD* mutations associated with exon-50 deletion in DMD therapy. Despite its high efficiency, this gRNA has a propensity to induce off-target editing in the *STRIP1* gene when used with SpCas9 (Ousterout et al., 2015) (Figure 5C). By using high-fidelity SpCas9 variants, we successfully mitigated the off-target editing. However, this came at the cost of reducing the editing efficiency by 9.1-17.9%, despite the utilisation of high-dose plasmid transfection followed by puromycin selection in our experiments. Notably, when we employed the 3TC scaffold for the high-fidelity SpCas9 variants, we successfully restored the on-target editing efficiency to levels comparable to those achieved with wild-type SpCas9 while still mitigating the off-target editing (Figure 5C). This suggests the benefits of employing the 3TC scaffold when utilising the high-fidelity SpCas9 variants, SpCas9-HF1 and eSpCas9(1.1).

### The 3TC Scaffold improves base editing

We then investigated whether the 3TC scaffold could enhance adenine base editing with SpCas9. Using the ABEmax system for A•T to G•C conversion (Koblan et al., 2018), we generated versatile all-in-one ABEmax constructs with either 4T or 3TC scaffolds. These constructs included a golden gate site for the easy insertion of spacer sequences and a puromycin-selection marker, akin to the PX459 system. Six different gRNAs targeting the mouse *Prl* locus, utilising either 4T or 3TC scaffolds were generated and tested in mouse embryonic stem cells. Our results demonstrated that the 3TC scaffold improved ABEmax efficiency, with four gRNAs utilising the 3TC scaffold exhibited significantly higher base editing efficiency compared to their 4T counterparts (Figure 6). Two gRNAs, Prl ABE-A and Prl ABE-F, demonstrated particularly remarkable improvements, elevating base editing efficiency from 44.2% to 92.9% and 77.3% to 99.6%, respectively. These results strongly support the use of the 3TC scaffold in base editing experiments to significantly optimize editing efficiency.

**Figure 6:**
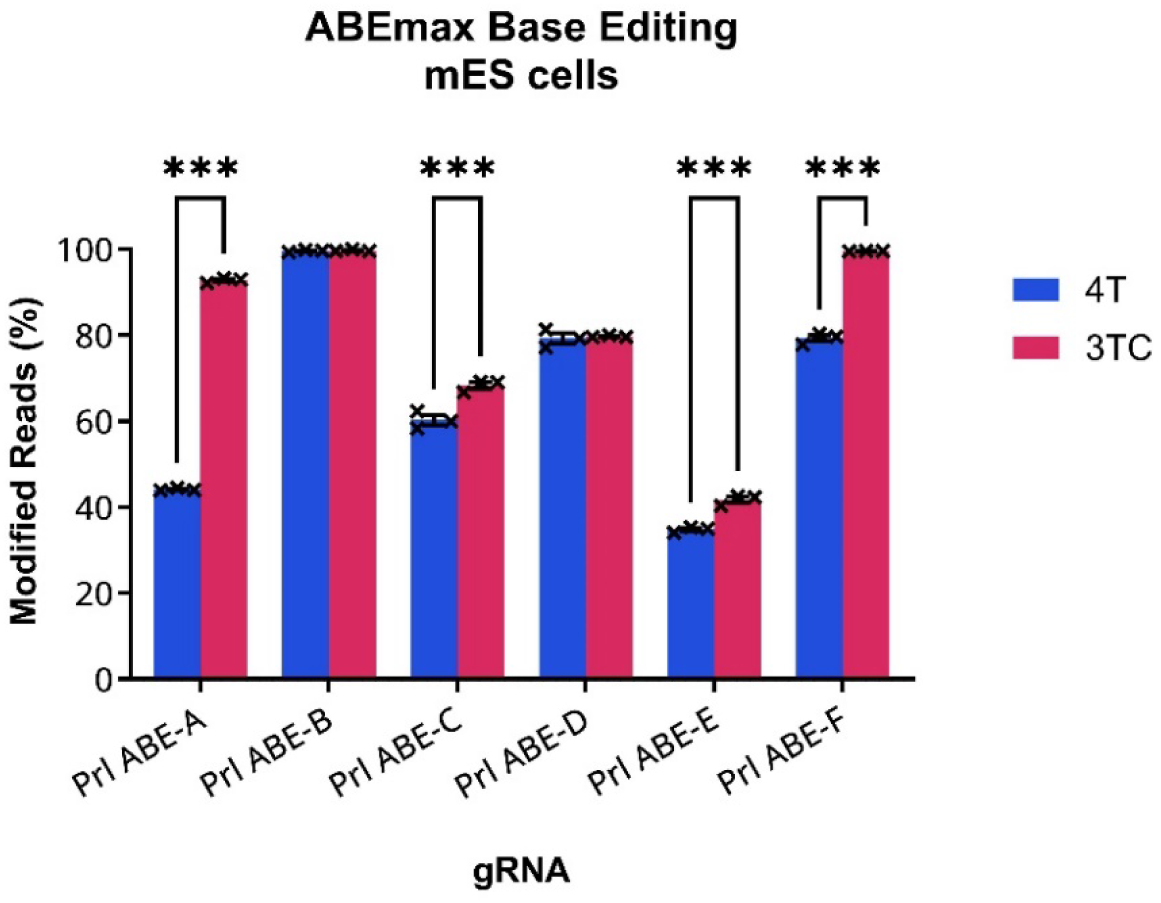
The 3TC scaffold improves ABEmax base editing. Base editing efficiencies of ABEmax delivered by lipofection of PX459v2-ABEmax (4T) and PX459v3-ABEmax (3TC) vectors to mES cells with 48-h puromycin selection, assessed by deep amplicon sequencing. Mean ± SEM; n=3. Two-way ANOVA with Šídák’s multiple comparisons test; ***p ≤ 0.001.

### The 3TC scaffold boosts SaCas9 gRNA transcript levels and editing efficiency

The SaCas9 gRNA scaffold can also undergo the 4T to 3TC modification, as it contains a four T-string in the gRNA scaffold (Figure 7A). We aimed to investigate whether this modification could enhance SaCas9 gRNA transcript levels and editing efficiency. To facilitate a direct and easy comparison between 4T and 3TC scaffolds, we constructed all-in-one SaCas9 plasmids with a golden gate site for easy gRNA cloning and a puromycin selectable marker, utilising either the 4T or 3TC SaCas9 gRNA scaffold. These plasmids are denoted as SaCas9-Puro.V2 and SaCas9-Puro.V3, respectively. We conducted a comparison of four different gRNAs targeting *DMD* in HEK293T cells to assess transcript levels and editing efficiency between SaCas9-Puro.V2 (4T scaffold) and SaCas9-Puro.V3 (3TC scaffold). As anticipated, we observed a significant increase in SaCas9 gRNA transcript expression (between 6-12 doublings) with the 3TC scaffold compared to the original 4T scaffold (Figure 7B-E). We then performed another independent transfection with high plasmid dose and puromycin selection to assess editing efficiency. Strikingly, the 3TC scaffold substantially enhanced overall SaCas9 editing (Figure 8A). For three gRNAs (hDMD-Sa-B, Sa-C, and Sa-D), the editing efficiency with the 4T scaffold was 44.1%, 3.6%, and 19.7%, and improved to 93.0%, 88.7%, and 94.2%, respectively, when utilising the 3TC scaffold. Another gRNA, hDMD-Sa-A, yielded similar editing efficiency with 4T or 3TC scaffold. Additionally, we tested five more gRNAs targeting *CXCR4*, previously assessed by Wang et al. (2017), and found that two gRNAs exhibited significantly higher editing efficiency, while the remaining three showed similar editing activity (Wang et al., 2017) (Figure 8A). Given our previous findings indicated that the 3TC scaffold improves editing under low vector availability, we tested the 4 gRNAs with similar activity between 4T and 3TC scaffold (hDMD-Sa-A, hCXCR4-Sa-g4, Sa-g9 and Sa-g12) under low vector conditions. We observed higher editing efficiency with the 3TC scaffold in three gRNAs (hDMD-Sa-A, hCXCR4-Sa-g4, and -g9), further confirming the utility of the 3TC gRNA scaffold modification for editing using SaCas9.

**Figure 7:**
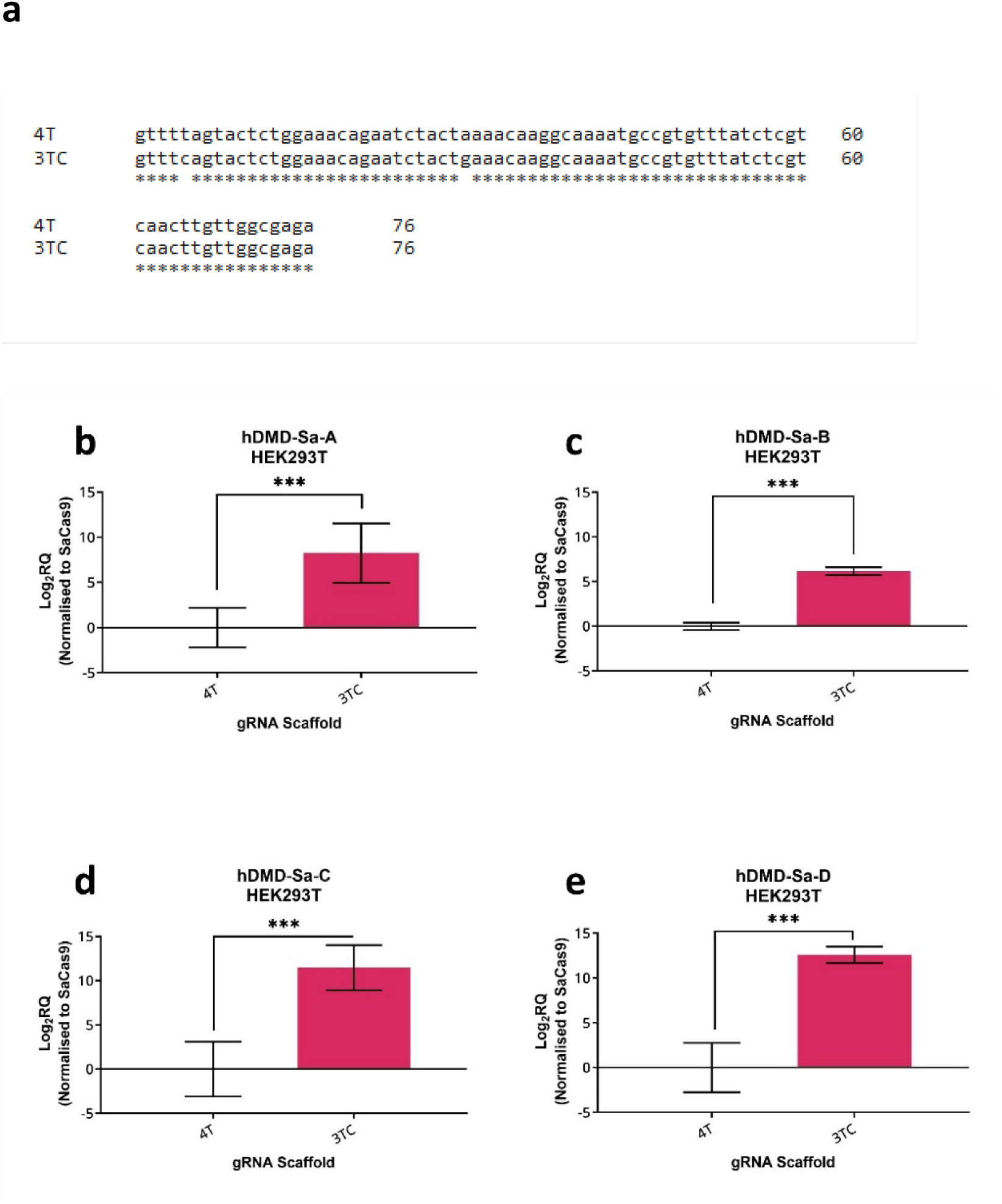
(A) DNA sequence of the 4T and modified 3TC scaffolds. (B-E) Sa gRNA expression levels are significantly increased with the modified scaffold. Relative quantification (RQ) of Sa gRNA expression levels of original scaffold SaCas9V2.Puro (4T) and modified scaffold SaCas9V3.Puro (3TC) plasmids delivered by nucleofection in HEK293T cells, measured by qRT-PCR. Data presented as mean Log2RQ normalised to SaCas9 mRNA expression ± 95% CI; n=3. Unpaired T-tests performed on ΔΔCT values; ***p ≤ 0.001.

**Figure 8:**
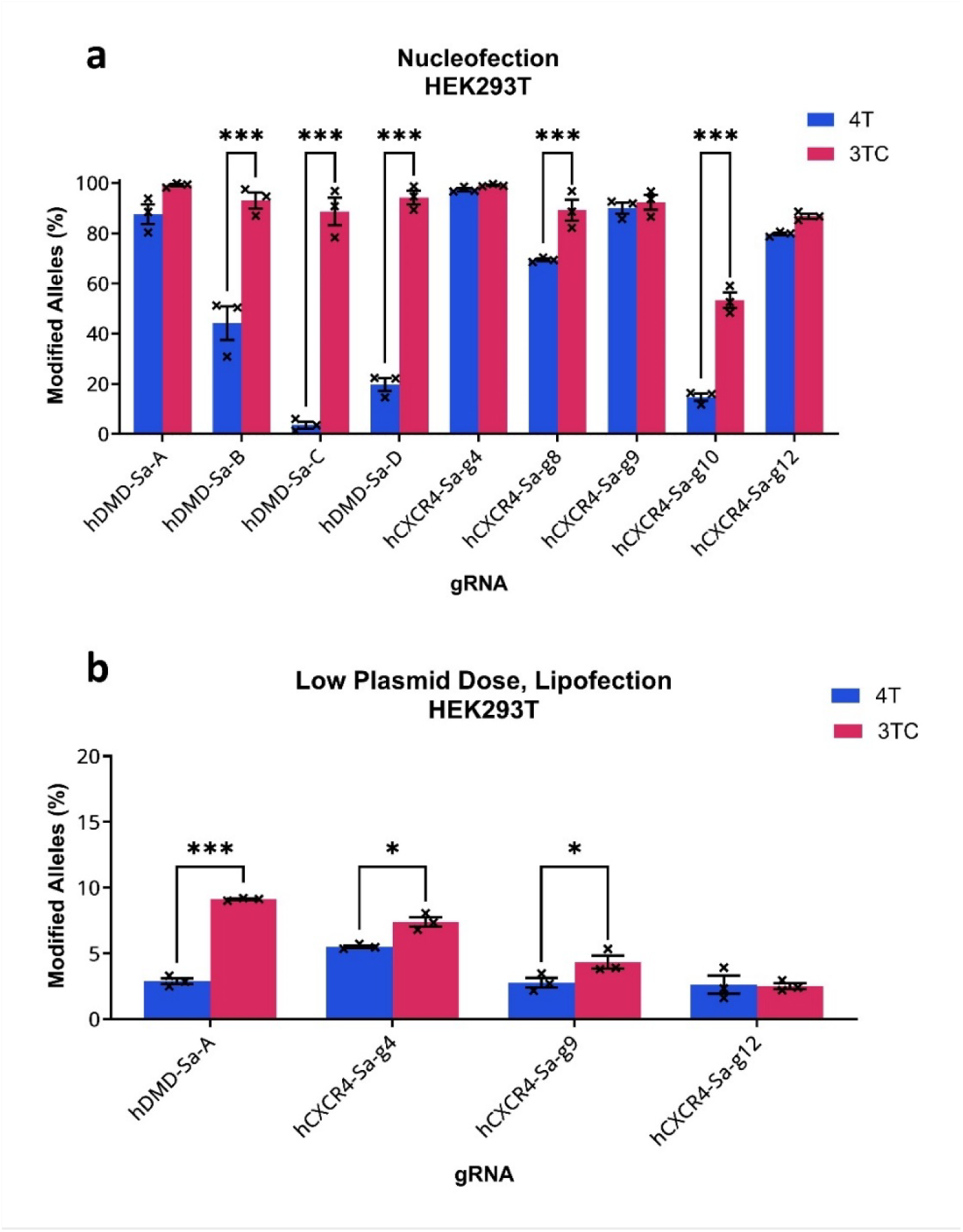
The 3TC scaffold improves SaCas9 editing efficiency. (A) Comparison of editing efficiencies of Sa gRNAs with original scaffold SaCas9V2.Puro (4T) and modified 3TC scaffold SaCas9V3.Puro plasmids in HEK293T cells delivered via nucleofection with subsequent puromycin selection, assessed by deep amplicon sequencing. Mean ± SEM; n=3; Two-way ANOVA with Šídák’s multiple comparisons test; **p ≤ 0.01, ***p ≤ 0.001. (B) Editing efficiencies of gRNAs delivered in low plasmid doses via lipofection without puromycin selection. Mean ± SEM; n=3; Two-way ANOVA with Šídák’s multiple comparisons test; *p ≤ 0.05, ***p ≤ 0.001.

To discern whether the enhancement of SaCas9 editing by the 3TC scaffold stems from increased transcript levels or intrinsic scaffold-induced activity, we conducted an in-vitro cutting assay. We compared gRNA hDMD-Sa-B with both the original and modified 3TC scaffolds produced by IVT. Employing a fixed gRNA amount, we observed no significant difference in SaCas9 cutting activity *in-vitro* (Supplementary Figure 1). This suggests that equivalent transcript levels from the 4T or 3TC scaffold yield similar cutting activity, implying that the elevated gRNA transcript levels facilitated by the 3TC scaffold likely contribute to the improvement of SaCas9 editing activity in cells.

### The 3TC scaffold improves SaCas9 editing activity for EDIT-101

Considering the enhancement of SaCas9 editing with the 3TC scaffold, we investigated whether this modification could increase the editing efficiency of the EDIT-101 trial therapeutic which employs SaCas9 with dual gRNAs. EDIT-101 (developed by Editas Medicine) is designed to treat the recessive IVS26 mutation in the *CEP290* gene, which causes a severe form of the inherited retinal dystrophy Leber congenital amaurosis (LCA) due to aberrant mRNA splicing (Leroy et al., 2021, Pierce et al., 2024). In EDIT-101, the SaCas9 and two gRNAs target the flanking intronic region of the IVS26 mutation. Deletion of the intervening region removes the novel splice donor site, restoring correct splicing (Maeder et al., 2019). We firstly generated dual-gRNA versions of the SaCas9-Puro.V2 and SaCas9-Puro.V3 plasmids, inserting a second U6-driven gRNA cassette that mimics the configuration of our previously published pDG459. We refer to these plasmids as pDG-SaCas9-Puro.V2 (4T scaffold) and pDG-SaCas9-Puro.V3 (3TC scaffold). Targeting constructs expressing the EDIT-101 gRNAs were generated using both plasmids and tested for intervening deletion efficiency in HEK293T cells. Deletion rates were determined by qPCR of gDNA, where the primers bind within the expected deletion and the amplicon contains the IVS26 mutation site. Thus, only alleles still harbouring the mutation site are amplified. We observed a significant enhancement in editing efficiency with the use of the 3TC scaffold, achieving a deletion rate of 32% compared to the 19% deletion rate observed with the 4T scaffold (Figure 9). This outcome underscores the potential of the 3TC scaffold to yield greater efficacy for EDIT-101.

**Figure 9:**
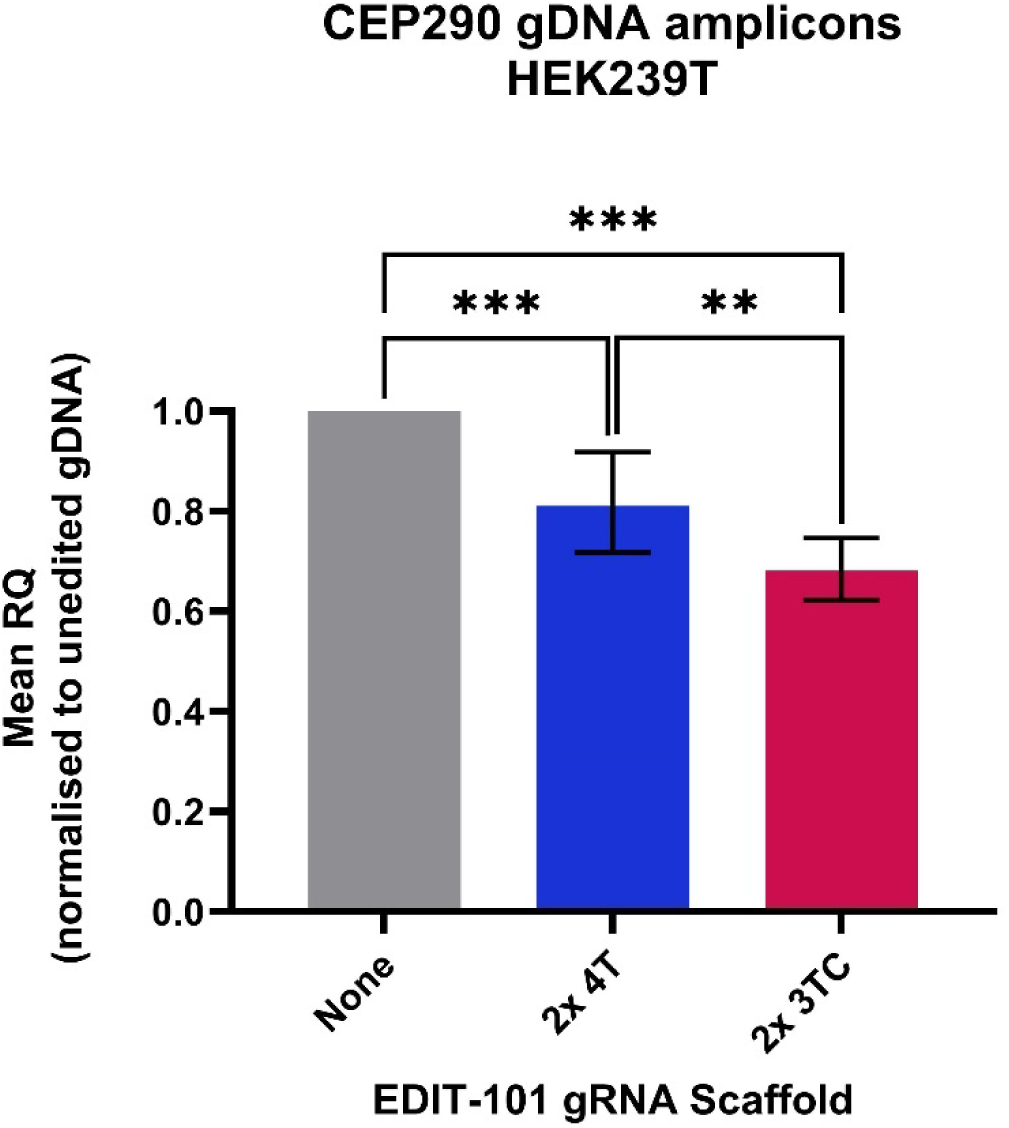
The 3TC scaffold improves SaCas9 editing efficiency in a dual-gRNA system compared to the original 4T scaffold in HEK293T cells. Unmodified *CEP290* gDNA alleles following delivery of pDG-SaCas9-Puro.V2 (2x4T) and pDG-SaCas9-Puro.V3 (2x3TC) by lipofection, measured by qPCR. The DMD gene was used as the endogenous unedited control for normalisation. Mean ± 95% CI, n=3 biological replicates. One-way ANOVA with Tukey’s multiple comparisons test performed on ΔΔCT values, **p ≤ 0.01, ***p ≤ 0.001.

## Discussion

Efficient genome editing using CRISPR-Cas9 gene editing technology is imperative for its successful application in a variety of research applications. Previous studies have underscored the pivotal role of gRNA transcript levels in determining editing activity (Jinek et al., 2013). In DNA-based CRISPR vectors, RNA Polymerase III (Pol III) is responsible for transcribing gRNAs under promoters such as U6, H1, or 7SK.

It has been observed that a string of four consecutive thymine (4T) residues serves as a minimal transcription terminator for Pol III (Arimbasseri and Maraia, 2015), capable of halting up to 75% of U6-driven transcription (Gao et al., 2018). Notably, the gRNA scaffold commonly utilised in SpCas9 or SaCas9 CRISPR systems contain this 4T string, which may contribute to suboptimal gRNA expression. Despite the presence of this suboptimal transcription terminator, these systems often exhibit high editing activity, suggesting that even with reduced gRNA transcript levels, there is still sufficient production for efficient gene editing. However, the specific nucleotide composition within the spacer sequence of the gRNA can directly influence its transcription efficiency. Notably, gRNA sequences rich in thymine residues (T-rich sequences) have been shown to exacerbate premature transcriptional termination, resulting in insufficient gRNA transcript production and consequently, lower gene editing efficiency (Graf et al., 2019, DeWeirdt et al., 2022).

We and other researchers have modified the string of consecutive thymine (T) bases from four (4T) to three (3T) in the gRNA scaffold to improve gRNA transcription (Chen et al., 2013, Deweirdt et al., 2020, Graf et al., 2019, Dang et al., 2015, Gao et al., 2019). In our study, we replaced the last thymine with cytosine (3TC) and its corresponding complement adenine (A) nucleotide with a guanine (G) nucleotide for both SpCas9 and SaCas9 gRNA scaffolds. Our findings demonstrate increased gRNA transcripts and improved editing outcomes for these nucleases. Additionally, we observed that this modification also enhances editing outcomes for high-fidelity SpCas9 nuclease variants and the SpCas9(D10A) adenine base editing system.

While the use of the 3TC scaffold does not uniformly lead to substantial enhancements, this could be attributed to the fact that gRNA expression levels with the 4T scaffold are already adequate for optimal editing in numerous instances. Therefore, increasing gRNA levels through the 3TC scaffold may not further enhance editing outcomes. However, there are scenarios where the use of the 3TC scaffold could offer advantages. For instance, when the combination of gRNA spacer sequences with the 4T scaffold results in insufficient gRNA transcripts, employing the 3TC scaffold could improve transcription levels, ensuring an adequate supply of gRNA transcripts for effective editing. Additionally, in cases where vector availability is limited, we demonstrated that gRNAs with the 3TC scaffold achieved higher editing efficiency compared to those with the 4T scaffold.

Given that in vivo CRISPR therapy often involves DNA vector delivery, such as AAVs, adenovirus, and lentivirus, where the dose is typically low for safety reasons or due to delivery limitations, we strongly recommend the adoption of the modified scaffold without the presence of the ≥4T string. This modification ensures optimal gRNA transcription and editing efficiency by eliminating sequences that signal a stop or pause for RNA Pol III. The 3TC scaffold could also be used for CRISPR library preparations and applications to maximise the activity of gRNAs used. We also propose that other Cas9 orthologs with a string of 4 Ts in the gRNA scaffold, such as *Streptococcus canis* Cas9 (ScCas9) and *Campylobacter jejuni* Cas9 (CjCas9) (Chatterjee et al., 2018, Kim et al., 2017), may potentially benefit from similar scaffold modifications.

## Materials and Methods

### Plasmid generation

PX459.V2 (Addgene #62988) was a gift from Feng Zhang (Ran et al., 2013). PX459.V2-SpCas9-HF1 (Addgene #108293) and PX459.V2 eSpCas9(1.1) (Addgene #108292) were gifts from Yuichiro Miyaoka (Kato-Inui et al., 2018). PDG459.V2 (Addgene #100901) was from our previously published study (Adikusuma et al., 2017). To generate the 3TC scaffold plasmids, these plasmids were modified by removing the U6-gRNA cassettes using PciI and Acc65I restriction digests and followed by ligation to insert a gBlock (IDT) fragment containing the cassette with the 3TC scaffold versions. SaCas9-Puro.V2 was generated by replacing the SpCas9 coding sequences of PX459.V2 with SaCas9 coding sequences from PX601 (Addgene #61591). Then the U6-(Sp)gRNA cassette was replaced with the U6-(Sa)gRNA fragment. To generate the SaCas9-Puro.V3, the U6-(Sa)gRNA cassette of SaCas9-Puro.V2 was replaced with the 3TC version. These plasmids are available through Addgene.

ABEmax (Addgene #112101) was a gift from David Liu (Koblan et al., 2018). PX459v2-ABEmax and PX459v3-ABEmax were generated by first digesting PX459v2 (Addgene #62988) and PX459v3 with AgeI (NEB) and PspOM1 (NEB) and gel purifying (Qiagen) the 7.7 kb fragment. The ABEmax construct (Addgene #112101) was amplified with primers ABEmax-F and ABEmax-R to enable digestion of the 2.6kb PCR product with AgeI (NEB) and PspOM1 (NEB) followed by PCR purification (Qiagen). These two fragments were ligated together and verified by whole plasmid sequencing (GNOMIX).

PdgSaCas9.V2 and V3 plasmids were generated by inserting a gBlock fragment containing an additional U6-driven gRNA cassette with either the 4T or 3TC scaffold into the XmaI and SbfI restriction site of SaCas9.V2 and SaCas9.V3, respectively.

Single gRNAs were inserted into backbone plasmids as previously described by Ran et al (2013). Briefly, the empty plasmids were digested with BbsI for 3 hours followed by gel electrophoresis separation. The linearized plasmids were then gel purified (Qiagen). Phospho-annealed oligo pairs carrying the guide sequences were ligated to the linearized empty plasmids. Plasmids were prepared using Qiagen miniprep kit and sequence validated by sanger sequencing. Multiple oligo insertion into dual-guide plasmids were performed using a previously described one-step cloning protocol (Adikusuma et al., 2017).

gRNA sequences and the corresponding oligo sequences are listed in Supplementary Information.

### Cell culture and transfections

HEK293T (ATCC CRL-3216) and C2C12 (ATCC CRL-1772) cells were maintained in DMEM (Gibco #11960044), supplemented with 10% fetal calf serum (FCS; Sigma) and 2 mM GlutaMax (Gibco). ES cells were maintained in DMEM, supplemented with 20% FCS, 2 mM GlutaMax, 100 µM non-essential amino acid, 100 µM 2-mercaptoethanol (Sigma), 3 µM CHIR99021, 1 µM PD0325901 and LIF. All cultures were kept in humidified incubators at 37°C and 5% CO_2_.

Nucleofections were performed using the Neon Transfection System 100 µL Kit (Invitrogen) following the manufacturer’s protocol, delivering 8 µg of plasmid DNA per transfection. HEK293T cells were nucleofected at 1.5 x 10^6^ cells/ 100 µL, at 1100 V, 20 ms and 2 pulses. Puromycin selection for HEK293T cells was applied 24 hours post-nucleofection at 2 µg/mL for 3 days. C2C12 cells were nucleofected at 1×10^6^ cells/ 100 µL, at 1400 V, 20 ms and 2 pulses. Puromycin selection for C2C12 cells was applied 24 hours post-nucleofection at 1 µg/mL for the first 24 hours and then 2 µg/mL for the subsequent 48 hours. mES cells were nucleofected at 1×10^6^ cells/ 100 µL at 1400 V, 10 ms and 3 pulses. Puromycin selection for mES cells was applied 24 hours post-nucleofection at 2 µg/mL for 48 hours. Following selection, cells were recovered to confluency in normal media without puromycin prior to collection for gDNA extraction.

For the lipofection experiments, HEK293T cells were seeded at 3×10^5^ cells per 6 cm dish. The next day, mixture of 10 µL of Lipofectamine-2000 (Invitrogen) and 3 µg (normal dose) or 10 ng (low dose) of plasmid DNA with OptiMEM was added to the dishes according to manufacturer’s instructions. 24 hours later, media was changed with fresh media and cells were harvested 3 days later. In the experiments with puromycin selection, 2 µg/mL of puromycin media was used during media change, and the puromycin media was refreshed daily for 3 days before harvesting. For EDIT-101 lipofections in HEK293T cells, 5 µL of Lipofectamine-2000 (Invitrogen) and 0.5 µg of plasmid DNA with OptiMEM was used. No puromycin selection was performed. Post-transfection, cells were grown to confluency for DNA extraction and analysis.

For ES cell lipofections, 6 µg of plasmid DNA and 6 µL of Lipofectamine-2000 (Invitrogen) were combined with OptiMEM (Gibco) with a final volume of 600 µL. This mixture was combined ES cells suspended in 100 µL of OptiMEM (Gibco) at a concentration of 0.75×10^6^ cells / 100 µL and incubated at 37°C for 15 minutes. These cells were seeded into T25 flasks containing warm ES cell media. 24 hours post-lipofection, cells were selected with puromycin at a concentration of 2 µg/mL for 48 hours. Following selection, cells were recovered to confluency in media without puromycin prior to collection for gDNA extraction.

### DNA extractions, PCR and NGS

gDNA was extracted from cell pellets using the High Pure PCR template preparation kit (Roche) as per manufacturer’s instructions. For deep amplicon sequencing, PCR was performed with Nextera-tagged primers. NGS was performed by Australia Genome Research Facility (AGRF) using the MiSeq Nano System, paired-end 500 cycle. Sequencing reads were analysed online using CRISPResso2 (Clement et al., 2019). Analysis was performed with these changes to default settings and quality filtering: Quantification window size of 5; Minimum average read quality >20; Minimum single bp quality of >10; Replace bases with N that have a quality lower than <10; Exclude 5 bp from the left side of the amplicon sequence for the quantification of the mutations.

PCR and NGS primer sequence lists are provided in Supplementary Information.

### q-RT PCR quantification

Quantification of SpCas9 gRNA transcripts was performed in C2C12 cells, while the SaCas9 gRNA was in HEK293T cells. C2C12 and HEK293T cells were nucleofected as described above except the number of cells per transfection was higher (2 x 10^6^ cells for HEK293T or 1.5 x 10^6^ cells for C2C12). High concentration of puromycin media (4 µg/mL) was added the following day. After 24 hours in the puromycin media, cells were collected for RNA extraction using the miRNeasy Mini kit (Qiagen) with DNase digest step as per manufacturer’s instructions. Reverse transcription was performed using SuperScript III Reverse Transcriptase (Invitrogen), using gRNA and Cas9-specific primers. Relative quantification of gRNA expression levels was performed by qRT-PCR and analysed by the ΔΔCT method, normalised to the Cas9 expression. qPCR was performed using Fast SYBR Green MasterMix Real-Time PCR Master Mix (Applied Biosystems) and cycled on the QuantStudio 3 Real-Time PCR System (Applied Biosystems).

Relative quantification of unmodified CEP290 gDNA alleles was performed by qPCR and analysed by the ΔΔCT method, normalised to unedited HEK293T DNA. qPCR was performed using Fast SYBR Green MasterMix Real-Time PCR Master Mix (Applied Biosystems) and cycled on the QuantStudio 3 Real-Time PCR System (Applied Biosystems). Prior to qPCR, gDNA was sheared by vortexing. gDNA from each sample was amplified in triplicate.

### In-vitro cleavage assay

gRNA was generated by in-vitro transcription using the HiScribe™ T7 In Vitro Transcription Kit (NEB) and purified using the RNEasy Mini Kit (Qiagen) as per manufacturer’s protocol. For each reaction, 30nM of sgRNA was pre-incubated for 10 min at 25°C with 30nM SaCas9-NLS (Biovision) in 1x Cas9 reaction buffer (Biovision). Following pre-incubation, 3nM of target DNA amplicon was mixed to each reaction and immediately incubated at 37°C. Reactions were sequentially stopped at 0.5, 1, 2 and 5 minutes by the addition of 0.8 units Proteinase K (NEB). Fragments were resolved on 1% agarose gel and band intensities were quantified on Image Lab (BioRad).

### Statistical analysis

Statistics were calculated on GraphPad Prism 10. Statistics for qPCR data were performed on ΔΔCT values.

## Acknowledgments

Research was performed at SAHMRI on Kaurna Country. We pay respects to the Kaurna people of the Adelaide Plains and to all Aboriginal and Torres Strait Islander people. SAHMRI is committed to embracing knowledge and culture as we continue our working journey to incorporate Aboriginal health research across all our themes and further reconciliation. The authors acknowledge the facilities and scientific and technical assistance of the cryogenics core facility at SAHMRI.

