## Supplementary Figure for "Enhancing gRNA Transcript levels by Reducing the Scaffold Poly-T Tract for Optimal SpCas9- and SaCas9-mediated Gene Editing"

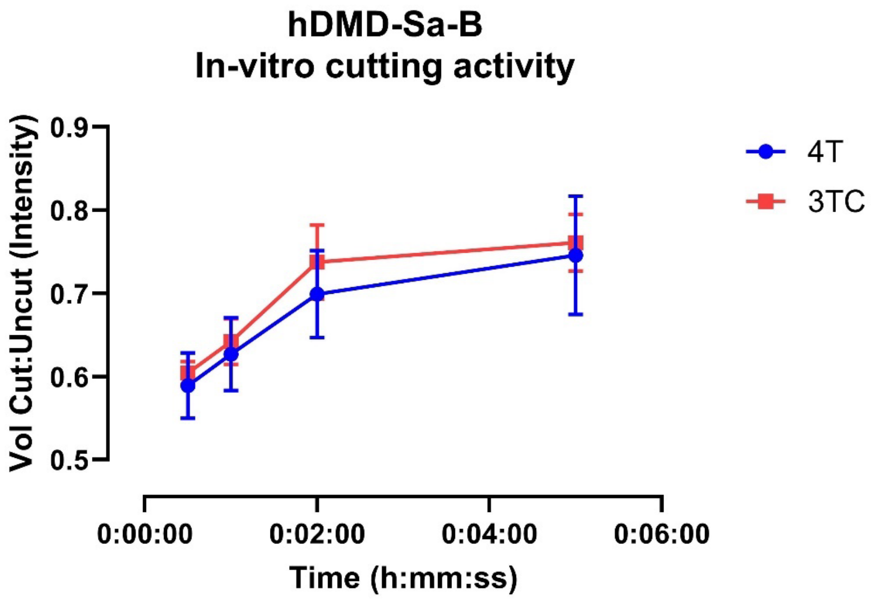


**Supplementary figure 1:** Band densitometry of the in-vitro cutting assay comparing the cutting rates of the hDMD Sa-B gRNA with either the 4T or 3TC scaffolds. Mean ± SEM.
